# Weakly alkaline conditions degrade aflatoxins through lactone ring opening

**DOI:** 10.1101/2023.10.31.564999

**Authors:** Natalie Sandlin, Jiu Lee, Marco Zaccaria, Marek Domin, Babak Momeni

## Abstract

Aflatoxins (AFs) are fungal metabolites that ubiquitously contaminate many common food crops and contribute to major foodborne diseases in humans and animals. The ability to remove AFs from common food and feed commodities will improve health standards and limit the economic impact inflicted by AF food contamination. Known chemical strategies have used strong acids and bases to remove contaminating AF, but these methods often lead to ecological waste issues downstream. In this study, we explore the application of weaker acidic and alkaline conditions to removes two types of AFs, AFB_1_ and AFG_2_. We find that a pH 9 buffered environment reduces AFB_1_ and AFG_2_ by more than 50% and 95%, respectively, within 24 hours. We show that AF degradation is through lactone ring opening, which is a known cause of AF toxicity, and provide a potential structure of the AFG_2_ degradation byproduct. Further, we confirm that incubation in the pH 9 environment reduces the genotoxicity of AFB_1_. Our findings indicate that a weakly alkaline environment may adequately detoxify AF-contaminated food or feed without the need to apply stronger or harsher basic conditions.

## Introduction

Aflatoxins (AFs) are fungal secondary metabolites produced by some *Aspergillus* species, which ubiquitously contaminate common food crops, such as corn, wheat, barley, and oats (1, 2). There are four major types of AFs: AFB_1_, AFB_2_, AFG_1_, and AFG_2_. The most common and toxic of the AFs is AFB_1_, which is classified as a Group 1 carcinogen and known to cause major health and economic crises worldwide. Consumption of AF-contaminated products leads to serious health effects in both animals and humans, including immunodepression, liver cancer, hormone disorders, and congenital malformation (3).

To prevent the harmful effects of AF contamination, strategies to remove them from food and feed must be implemented. Current decontamination strategies are classified as physical, chemical, and biological methods. Physical methods include sorting, milling, washing, and irradiation steps that have low reproducibility and high cost (4–6); conversely, biological methods consist of using living organisms and/or their products (e.g. microbial enzymes) to remove the toxins, where limitations lie in accurate determination of mechanisms and optimal working conditions (7, 8). Alternatively, current chemical methods involve the conversion of AF through applications such as ammonization, ozonation, and peroxidation (5, 9).

A known chemical strategy for AF removal is through the application of strong alkaline or acidic conditions (4, 10). For example, Mendez-Albores *et al*. found that the addition of 1 N aqueous citric acid converted AFB_1_ into less toxic byproducts (11). Alternatively, KOH treatment (pH 12) by Vidal *et al*. significantly reduced AF to below the limits of detection (12). Additionally, in the context of alkaline conditions to eliminate AF from foodstuffs, there are current treatment practices such as alkaline electrolyzed water, alkaline cooking, and nixtamalization (13–15). Nixtamalization is a common and long-used processing technique used in the preparation of masa from corn. This process involves cooking the corn in boiling water containing lime (Ca(OH)_2_) to alkalinize the environment to a pH higher than 10 (16). While some chemical processing methods can reduce the availability of nutrients in food, the process of nixtamalization can actually increase them. Nixtamalization effectively reduces the levels of AF as well as prepares the maize for downstream use; however, on the industrial scale, it produces large quantities of wastewater, polluted with organic matter and high in pH, that is difficult to dispose of (15). These examples support the removal AF contamination from foods employing strong alkaline and acidic conditions. However, to deploy these methods, we need to ensure that the chemicals are fully removed after processing, the nutritional value of the food is not diminished, and the ecological waste is kept to a minimum. These are the factors that limit the use of strong acids or alkalis to remove contamination in foods.

In this study, we consider the effects of weaker acidic and alkaline environments on the elimination of AFs, particularly AFB_1_ and AFG_2_. We test the degradation levels of buffered medium in the range of pH 4.0-9.0 using the native fluorescence of AFs and mass spectrometry as detection methods. The fluorescence of AF is strongly linked to its lactone ring moiety, also the main actor in AF’s toxicity (17). It has been discovered that opening of the lactone ring significantly reduces toxicity and fluorescence, making this a reliable proxy for AF detoxification (17). Further, we consider the degradation products and toxicity of pH 9 buffered medium as we propose a weakly alkaline environment as a means to remove AF from foods during processing.

## Results

### Increasing the pH of the medium leads to loss of aflatoxin fluorescence

We first quantified how different pH values affected the fluorescence of AF (see *fluorescence degradation assay* in Methods). A standard defined culture medium was buffered to pH 4-6 in 0.1 M citrate buffer, 7-8 in 0.1 M MOPS, and 8-9 in 0.1 M Tris-HCl. Buffered medium was then supplemented with AFB_1_ or AFG_2_ (at an initial concentration of 15 μg/mL) and degradation measured using our fluorescence assay. The fluorescence of AF in these buffered media was monitored during 48 hours of incubation time. The results for the full range of pH tested are shown in Fig. S1. Here, we explore the extremes of the pH range as well as a neutral pH for comparison. A neutral pH of 7 had little effect on the fluorescence of AFB_1_, while the extremes of the pH range showed increased fluorescence at acidic conditions and decreased fluorescence at basic conditions (Fig. 1). Particularly, pH 4 buffered medium displayed a rapid increase in the fluorescence of AFB_1_, but not AFG_2_ (Fig. 1). However, pH 9 buffered medium decreased fluorescence by ∼50% for AFB_1_ and by ∼95% for AFG_2_ (Fig. 1), suggesting a loss of toxin concentration. Additionally, to ensure that the effect was independent of the buffer used in the experiment, medium buffered to pH 9 using 4 different buffers was tested and showed consistent results of decreased AF fluorescence over the testing period (Fig. S2).

**Figure 1:**
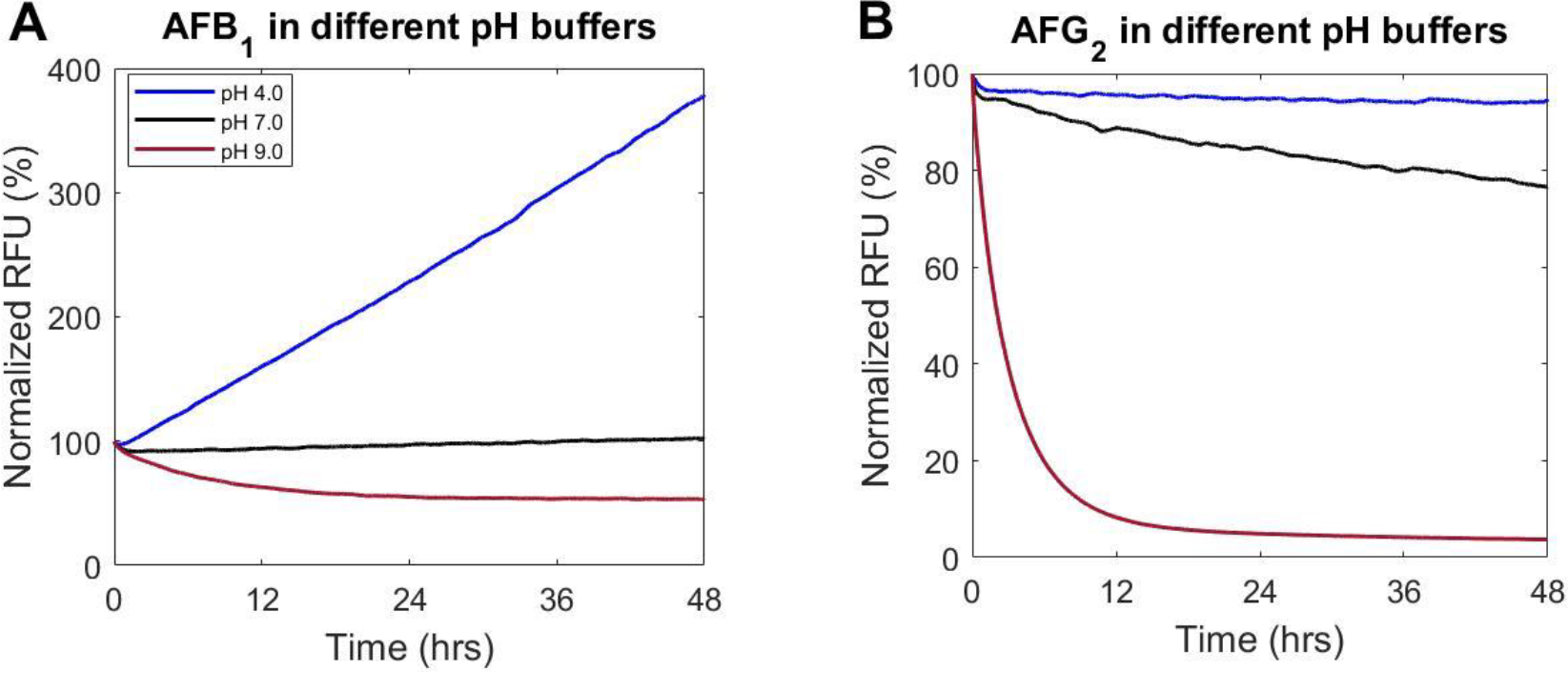
The pH in a buffered medium influences fluorescence readout in AF degradation assay. A) AFB_1_ and B) AFG_2_ were incubated with buffered medium at three pHs (4.0, 7.0, and 9.0) over 48 hours with readings for fluorescence of AF taken periodically over the incubation period. pH 4.0 was buffered in 0.1 M citrate buffer, pH 7.0 was buffered in 0.1 M MOPS, and pH 9.0 was buffered in 0.1 M Tris-HCl. Data is the mean of 3 replicates. RFU has been normalized to initial fluorescence.

### Loss of fluorescence by pH 9 medium is proportional to loss of toxin concentration

To test if the trends seen via our fluorescence assay represented AF degradation, we examined the changes in AF concentrations (initial concentrations set at 15 μg/mL) for pH 4 and pH 9 using an extraction/fluorescence method and LC-MS. The extraction method (see *aflatoxin extraction* in Methods) uses an ethyl acetate liquid/liquid extraction protocol to remove AF from the surrounding aqueous environment. Such extraction essentially eliminates any transient effects of the pH on the toxin. Pairing this extraction with fluorescence analysis allows for an accurate reading of AF levels independent of the short-term effects of the pH environment. After extraction at three time points, 0, 24, and 48 hours, pH 4 samples showed insignificant changes to the AF levels for both AFB_1_ and AFG_2_ (Figs. 2A and 2B), indicating that the increases to the fluorescence in the initial tests (Fig. 1) were transient effects of the pH on AFs. In contrast, pH 9 samples displayed sustained degradation of AFB_1_ and AFG_2_ by ∼50% and >95%, respectively, within 24 hours (Figs. 2A and 2B). Additionally, an LC-MS analysis of the pH 9 conditions (see *LC-MS assay* in Methods) showed similar results in the remaining levels of AFB_1_ (Fig. 2C) and AFG_2_ (Fig. 2D) after 48 hours incubation, similar to the extraction/fluorescence method of detection. First, this further confirms that pH 9 buffered medium degrades AFs. Second, these two methods taken together show that the loss of AF fluorescence in pH 9 conditions is associated with a decrease in the toxin concentration.

**Figure 2:**
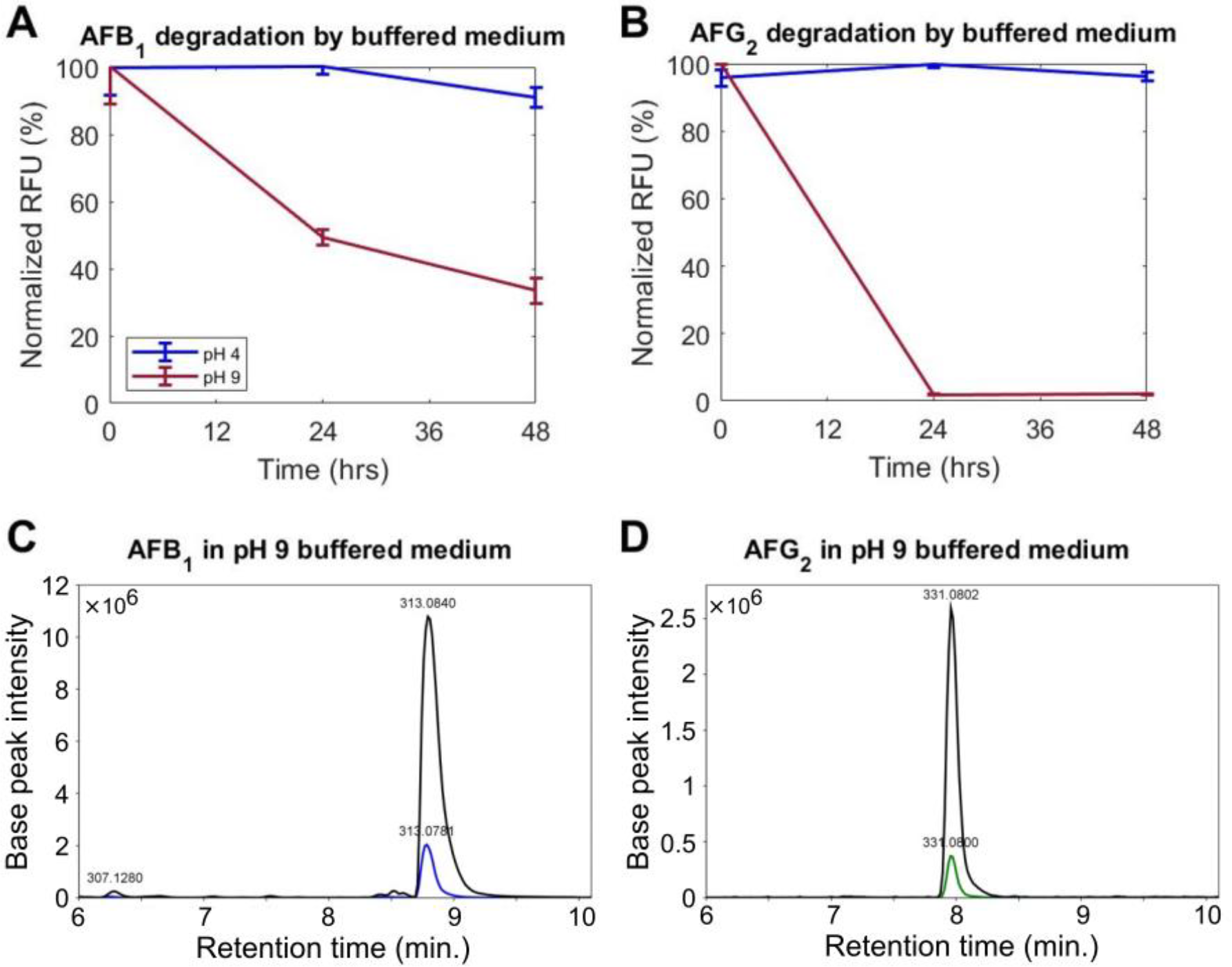
pH 9 buffered medium degrades AFs. Normalized fluorescence of A) AFB_1_ and B) AFG_2_ from ethyl acetate extraction after incubation buffered medium at time points (TP) 0, 24, and 48 hours, shows a decrease at pH 9, but not at pH 4. Data are the mean of three replicates. Error-bar shows the standard deviation. Sample analysis through LC-MS of C) AFB_1_ and D) AFG_2_ levels at TP 0 and 48 hours of incubation in pH 9 buffered medium corroborates the extraction/fluorescence results. The corresponding *m/z* value is shown for each identified peak.

### Degradation of AFs in a pH 9 buffered condition leads to lactone ring opening

Since the pH 9 buffered medium displays a promising AF degradation, we investigated potential byproducts of the degradation reaction. Based on the loss of fluorescence after incubation in pH 9 buffered medium and because of the link between the lactone ring and the fluorescence of AF (17), we hypothesized that the lactone ring moiety to be broken open. The LC-MS assay finds that as AFG_2_ in the sample is removed over the 48-hour incubation period, a second peak in the spectrum increases at m/z 305. Using the exact mass, a potential structure of the degradation byproduct was produced with opening of the lactone ring and a decarboxylation event (Fig. 3). A byproduct peak is not found in the AFB_1_ spectra, despite the decrease in toxin after incubation in the pH 9 buffered medium (Fig. S4). This indicates that AFB_1_ is likely degraded to a further extent than what is detectable through our analysis. However, due to the loss of fluorescence in AFB_1_ after incubation, we can infer that the lactone ring was targeted in a fashion similar to AFG_2_.

**Figure 3:**
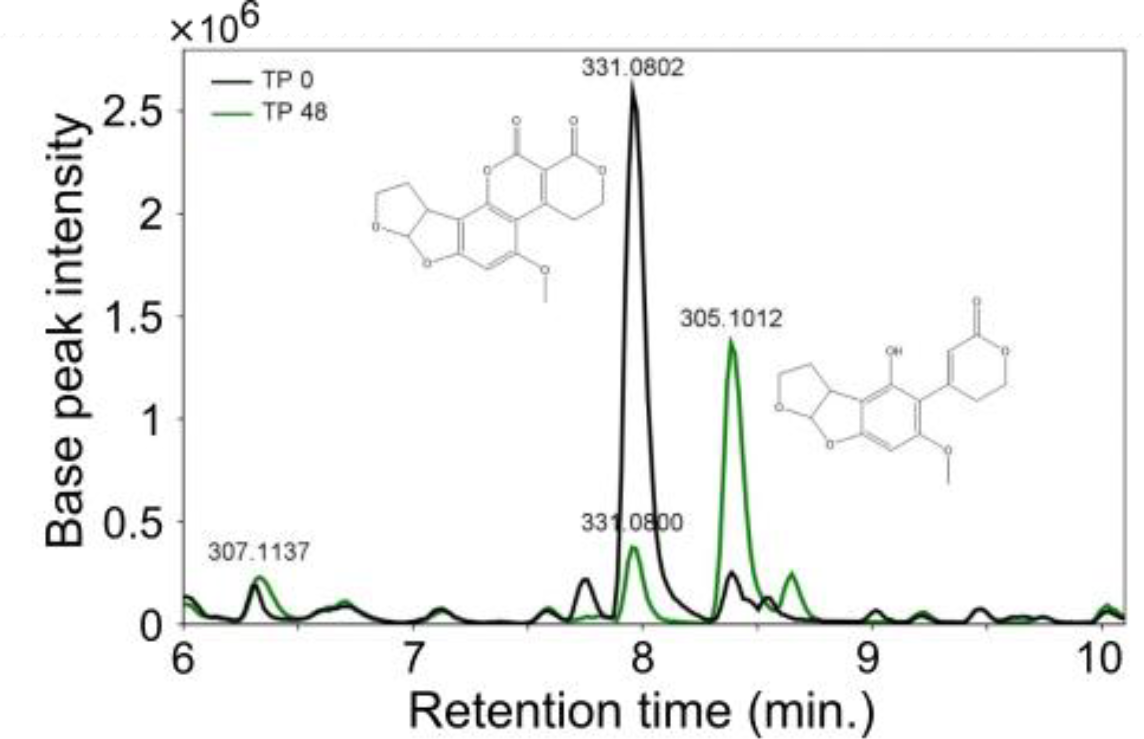
LC-MS analysis of AFG_2_ after incubation in pH 9 buffered medium reveals potential byproducts of degradation. Mass spectra for time points (TP) 0 and 48 hours of incubation of AFG_2_ in pH 9 buffered medium, highlighting *m/z* 331 (AFG_2_) and *m/z* 305 (potential byproduct). The corresponding *m/z* value is shown for each identified peak. Structures of AFG_2_ and the proposed structure of the byproduct are shown next to their respective peak.

### Byproducts of degradation in a pH 9 buffered condition have reduced toxicity

While it is confirmed that AF levels were decreased or diminished in the pH 9 buffered condition, this alone does not confirm that toxicity has also been decreased. To test the toxicity of the byproducts of degradation in this condition, we conducted an SOS ChromoTest assay for genotoxicity, using the S9 rat liver enzyme induction (see *genotoxicity assay* in Methods). This emulates the post-consumption induction by cytochrome P450 that activates AF toxicity (18). Aflatoxin B_1_ was incubated in the pH 9 condition for 48 hours prior to testing. We used controls of toxin levels in a neutral pH medium (pH 7) that did not show degradation, according to our fluorescence assay (Fig. 1) and LC-MS analysis (Fig. S3). Standard interpretation of this assay considers an IF of >1.5 as genotoxic. In our results, the AFB_1_ control condition had an IF of ∼2.6 at the highest concentration tested (15 μg/mL), therefore indicating genotoxicity. The same starting concentration of AFB_1_ after incubation in the pH 9 condition reduced the IF to below 1.5, suggesting decreased toxicity (Fig. 4). Compared to controls, the pH 9 treated samples had reduced toxicity. Likely, the remaining toxicity in this sample came from the residual, undegraded AFB_1_ seen in fluorescence and MS experiments and not the byproducts of the degradation reaction. The toxicity of AFG_2_ byproducts was not investigated since the SOS ChromoTest is not sensitive for AFG_2_, but byproducts of the pH 9 condition showed loss of fluorescence and lactone ring opening, which have been confirmed to correlate to reduced toxicity (17).

**Figure 4:**
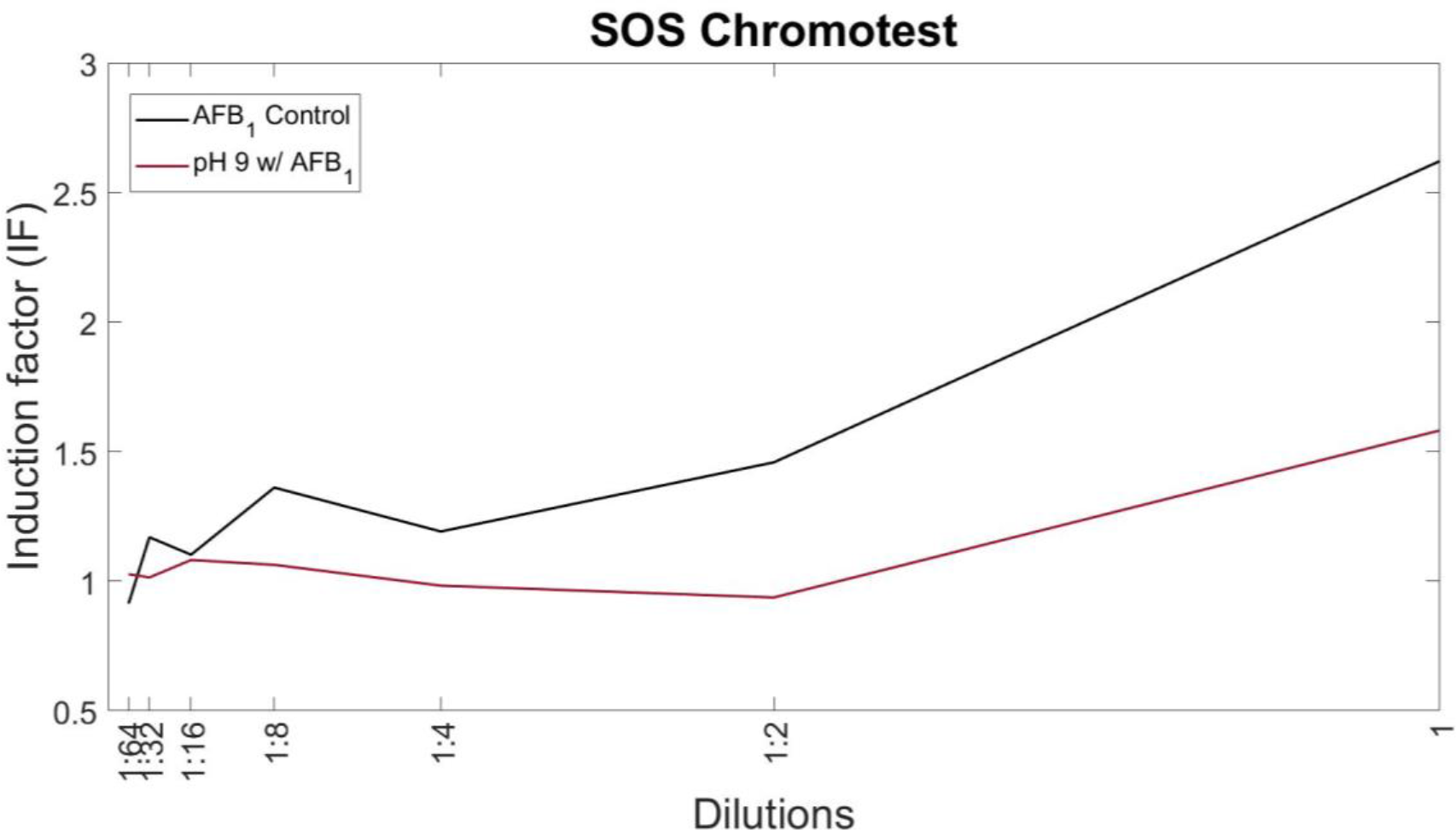
Byproducts of AFB_1_ degradation in pH 9 buffered medium have reduced toxicity. SOS-ChromoTest kit for genotoxicity was used to assess the toxicity of byproducts after pH 9-mediated degradation. AFB_1_ was incubated in pH 9 buffered medium for 48 hours prior to ethyl acetate extraction and resuspension in methanol. Serial 2-fold dilutions were used to obtain a trend line. AFB_1_ incubated in pH 7 medium was used as a control (no degradation) and run through the same extraction protocol. Values of IF over 1.5 are interpreted as genotoxic.

## Discussion

We investigated the ability to chemically detoxify prevalent crop contaminants, AFB_1_ and AFG_2_, through the application of weakly basic and acidic buffered medium conditions. We found that alkaline environments have the potential for AF degradation, consistent with previous literature (19, 20). A pH 9 buffered environment displayed degradation in our fluorescence assay to levels of 60% for AFB_1_ and >95% for AFG_2_ within 48 hours. We showed that the buffer used for pH 9 conditions does not significantly impact the degradation efficiency. Additionally, to our knowledge, we are the first to identify and test toxicity of degradation byproducts from alkaline conditions for AF degradation.

From our findings, we propose that the alkaline environment enables the degradation of AFs through the mechanism of lactone ring opening, potentially a spontaneous hydrolysis event. Alkaline hydrolysis is consistent with the byproducts of AF conversion in pH 9 conditions, in our study. Other microbe-produced molecules with lactone rings, such as quorum sensing molecules, are also known to degrade in alkaline environments (21, 22). However, to formally confirm the mechanism, further experimentation needs to be performed.

While our findings highlight the potential of pH manipulation as a method to degrade AFs, it is important to consider the practical implications and limitations of this approach. Chemical methods of AF decontamination have been widely studied and used in the agricultural industry. Yet, they are still limited by the arduous downstream clean up after treatment, especially full removal of chemicals and reduced ecological waste. Also, pH adjustment may not always be feasible in certain food processing or storage conditions, as it can impact the properties of the product or interfere with other desired chemical reactions. Previously used methods that implement alkaline environments have particularly struggled with the waste produced due to the strong bases used at large scales. However, these previous methods established a building block for our proposed strategy. Based on other applications of alkalis to remove AFs, this method of pH 9 buffered medium could be readily implemented in food processing at the stages of food washing (23). With the opening of the lactone ring by the alkaline conditions, the compound can become more water soluble and the AFs can be removed during washing with water. Since the pH of the method outlined in this study is less harsh than previous alkaline applications, we foresee water waste and other potential ecological pollutants can be reduced.

Another consideration is combining chemical alkaline methods with emerging biological means for decontamination. It is known that a number of bacterial and fungal species are capable of degrading AFs, for example by enzymatic conversion of AFs to non-toxic byproducts (24–26). Some of these species may turn the environment more basic, or achieve higher efficiency in a more alkaline environment. An interesting further study may be in the pursuit of a combined degradation effect utilizing an alkaline medium that supports the growth of such biological degraders to produce an enhanced degradation of AFs.

As a word of caution, we encourage future studies to use a neutral pH buffered environment and document the pH when screening for biological organisms that are capable of degrading AFs. This is critical for distinguishing the enzymatic degradation from alkaline degradation of AFs by different organisms. The absence of information about the pH in some of the previous reports of AF degradation makes the interpretation of the degradation mechanism difficult.

In conclusion, our study demonstrates that pH manipulation can be an effective strategy for the degradation of AFs. Alkaline conditions promote the hydrolysis of AFs, as seen in the proposed degradation product, and render them more susceptible to degradation mechanisms. The efficacy of pH-based degradation may vary depending on the AF type as seen in the differences between AFB_1_ and AFG_2_. Further research is needed to optimize pH manipulation approaches for different food and feed commodities and to evaluate their feasibility and practicality. The findings from this study contribute to our understanding of AF degradation mechanisms and offer insights for the development of effective strategies to reduce AF contamination in the food and feed industries.

## Methods

### Reagents

AFB_1_ and AFG_2_ (Cayman Chemical) were dissolved in LC-MS grade methanol to the final concentration of 1 mg/mL for stock solutions.

### Medium and pH buffering

Minimal medium was used as the base for all pH testing: KH_2_PO_4_ (1.5 g/L), K_2_HPO_4_ x 3H_2_O (3.8 g/L), (NH_4_)_2_SO_4_ (1.3 g/L), sodium citrate dihydrate (3.0g/L), FeSO_4_ (1.1 mg/L), 100x vitamin solution (1 mL), 1000x trace elements solution (1 mL), 1 M MgCl_2_ (5 mL), 1 M CaCl_2_ (1 mL), 100x amino acid stock (10 mL), and glucose (4.0 g/L). Stock solutions of citrate buffer, MOPS, and Tris-HCl were made at a concentration of 1 M prior to addition to the base minimal medium at a final concentration of 0.1 M.

### Aflatoxin extraction

For each indicated time point, liquid-liquid extraction was used to stop the reaction and extract aflatoxin and potential byproducts. Ethyl acetate was added (v/v) to each sample in microcentrifuge tubes and vortexed vigorously for 2 minutes. Samples were centrifuged at 11000 ×g for 2 minutes to separate the phases. The top aqueous layer was transferred to a new microcentrifuge tube. This extraction was repeated twice per sample to increase efficiency. Ethyl acetate was left to evaporate at 65°C. The aflatoxin precipitate was resuspended in 100-200 μL of LC/MS grade methanol.

### LC-MS assay

For the analysis we employed a Kinetex 2.6 μm EVO C18 column (100 × 2.1 mm). Mobile phase A: Water 5 mM Ammonium Acetate, 0.5% Acetic Acid. Mobile phase B: Methanol 5 mM Ammonium Acetate, 0.5% Acetic Acid. Flow rate was 350 μL/min. UV detection wavelength was set at 354, 360 nm. The following gradient method was used in all runs:

| Time (min) | %A | %B |
| --- | --- | --- |
| Initial | 90 | 10 |
| 3 | 90 | 10 |
| 10 | 30 | 70 |
| 10.1 | 10 | 90 |
| 12 | 10 | 90 |
| 12.1 | 90 | 10 |
| 15 | 90 | 10 |

The eluent from the column was directed into the electrospray source of an Agilent 6230 TOF mass spectrometer operated in positive ionization mode. Data was converted into the mzML file format and analyzed using the MZMine software.

### Fluorescence degradation assay

Buffered medium was aliquoted into sterile microcentrifuge tubes and aflatoxin was added according to desired final concentration (typically 15 μg/mL) per well. Samples were arrayed in black glass-bottom 96-well plates (Nunc™ #165305 96-Well Optical Bottom) at a final volume of 150 μL per well. Standard control of no toxin (medium alone) was used. A BioTek Synergy Mx multi-mode microplate reader was used to monitor optical density at 600 nm (to account for contamination) and fluorescence of aflatoxin at an excitation of 380 nm and emission of 440 nm with a gain of 50 for AFG_2_ and a gain of 65 for AFB_1_. Reads were taken at 5 min intervals over 48 hours (unless otherwise noted). Output was exported as a text file for downstream analysis and visualization. Typically, 2-3 replicates were used per condition. Sterile water was placed at the peripheral wells of the 96-well plate to contain evaporation.

For experiments testing fluorescence after extraction, a similar protocol was followed but with single point reads taken in triplicate.

### Genotoxicity assay

The genotoxicity assay was performed, with metabolic activation using S9 rat liver extract, according to the protocol for SOS-ChromoTest^TM^ kit from Environmental Bio-detection Products (Mississauga, ON, Canada). For the assay, the negative control was composed of a 10% dimethyl sulfoxide (DMSO) solution in sterile water, and the positive control was 2-aminoanthracene (2-AA). Toxin samples were prepared by adding AFB_1_ to pH 9 buffered medium at a concentration of 15 μg/mL and incubating at 28°C for 48 hours. A control of AFB_1_ in a pH 7 buffered medium (no degradation) incubated under the same conditions was used. After incubation, samples underwent extraction via the above described method prior to serial 2-fold dilutions, per the assay protocol. Kit-provided bacterial suspension (*E. coli*) was added to the toxin samples and incubated at 37°C for 2 hours with shaking. Color development was quantified on a BioTek Synergy Mx multi-mode microplate reader with readings at 420 nm to measure cell survival through alkaline phosphatase and at 600 nm to measure SOS system induction via β-galactosidase activity. SOS-induction factor (IF) was then calculated according to the provided analysis protocol.

## Author contributions

Conceptualization, N.S. and B.M.; methodology, N.S., J.L., M.D., and M.Z.; formal analysis, N.S.; investigation, N.S., M.D., and B.M.; data curation, N.S.; writing—original draft preparation, N.S.; writing—review and editing, N.S., M.Z., and B.M.; visualization, N.S.; supervision, B.M.; funding acquisition, N.S. and B.M. All authors have read and agreed to the published version of the manuscript.

## Data availability

The data that support the findings of this study are openly available at https://github.com/nsandlin7/pH_aflatoxin. Mass spectrometry files presented in this work will be available from the authors upon request.

## Acknowledgments

This work was partially supported by the National Science Foundation (NSF-CBET) under Grant No. 2103545. NS is supported by a NIFA-AFRI Predoctoral Fellowship from the USDA (Award No. 2021-67034-35108). We thank Marek Domin and the Boston College Mass Spectroscopy Core for infrastructure and support.

## Supplementary figures

**Figure S1.**
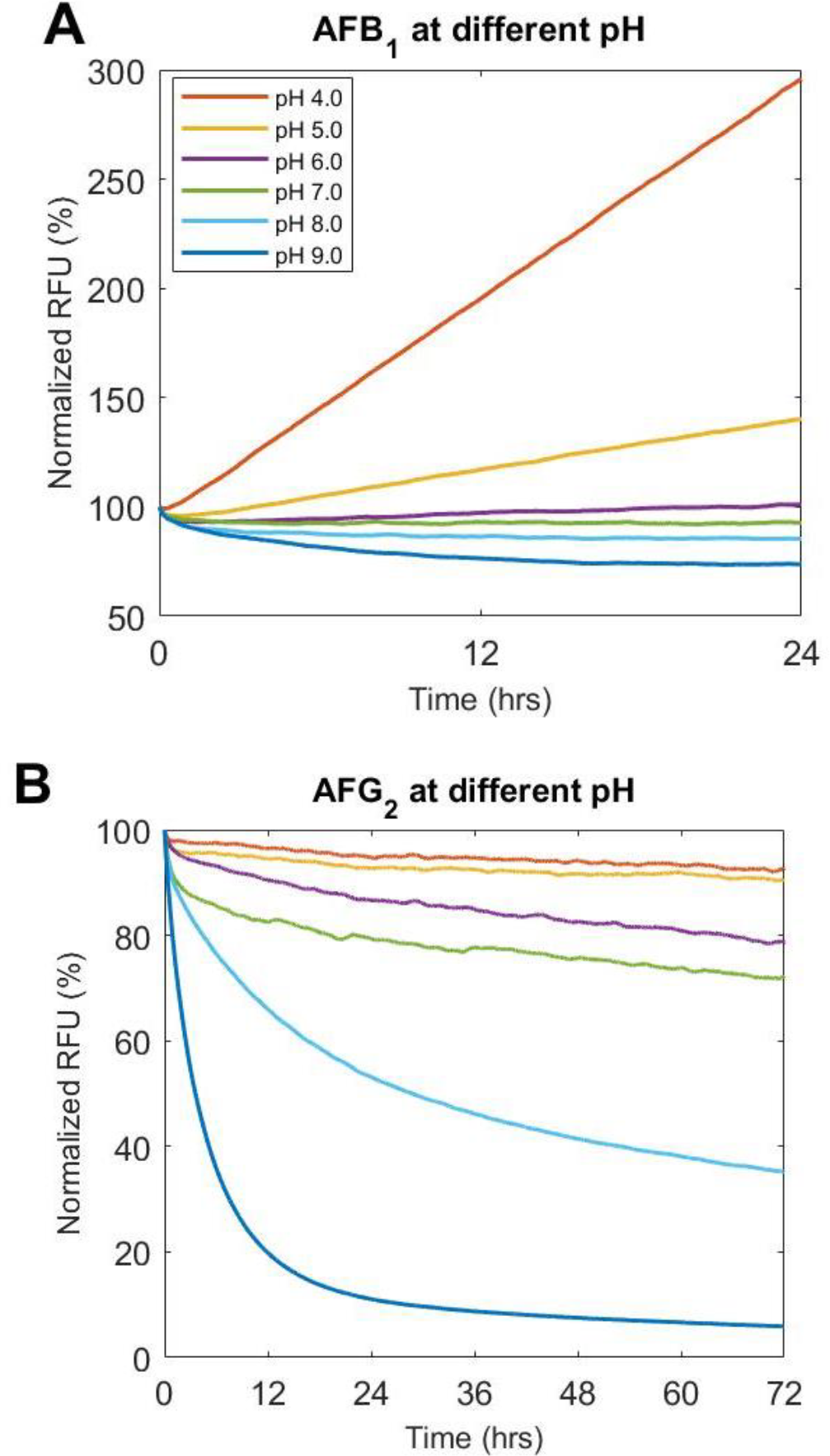
The fluorescence of AFB_1_ and AFG_2_ decreases in alkaline pH buffered medium. A) AFB_1_ and B) AFG_2_ were incubated with buffered medium at pHs within the range of 4.0 to 9.0 over 24 or 72 hours with readings for fluorescence of AF taken periodically over the incubation period. pH 4.0-6.0 was buffered in 0.1 M citrate buffer, pH 7.0 was buffered in 0.1 M MOPS, and pH 8.0-9.0 was buffered in 0.1 M Tris-HCl. RFU has been normalized to initial fluorescence.

**Figure S2.**
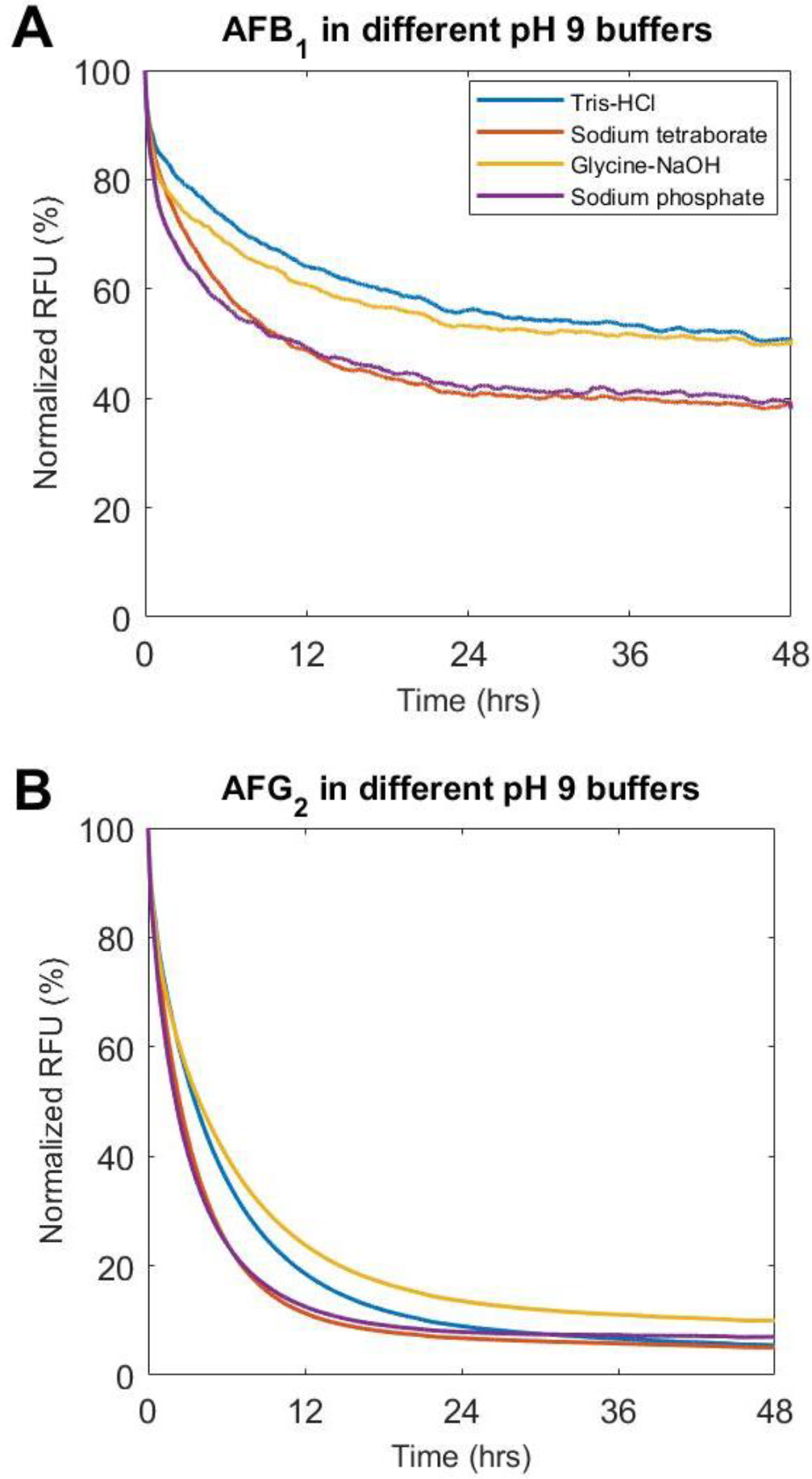
Different buffers affect AFB_1_ and AFG_2_ fluorescence similarly at pH 9. A) AFB_1_ and B) AFG_2_ were incubated with medium buffered at pH 9 in either 0.1 M Tris-HCl, sodium tetraborate, glycine-NaOH, or sodium phosphate for 48 hours with readings for fluorescence of AF taken periodically over the incubation period. RFU has been normalized to initial fluorescence.

**Figure S3.**
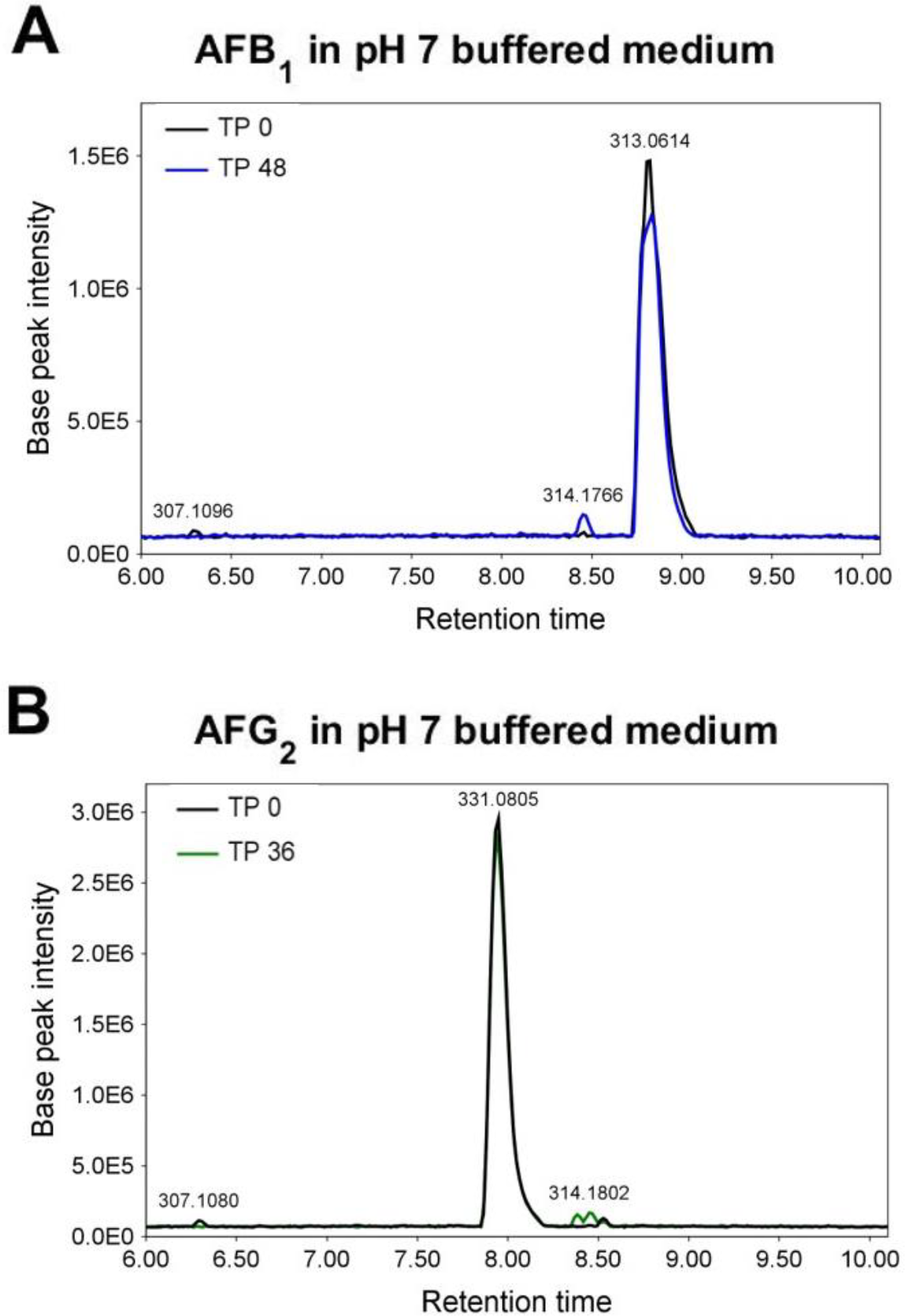
Neutral pH of 7 does not significantly affect aflatoxin levels. LC-MS analysis of A) AFB_1_ and B) AFG_2_ levels at time points (TP) 0 and 48 hours of incubation with pH 7 buffered medium shows that the AF concentrations are maintained over time. Retention times are in minutes. The corresponding *m/z* value is shown for each identified peak.

**Figure S4.**
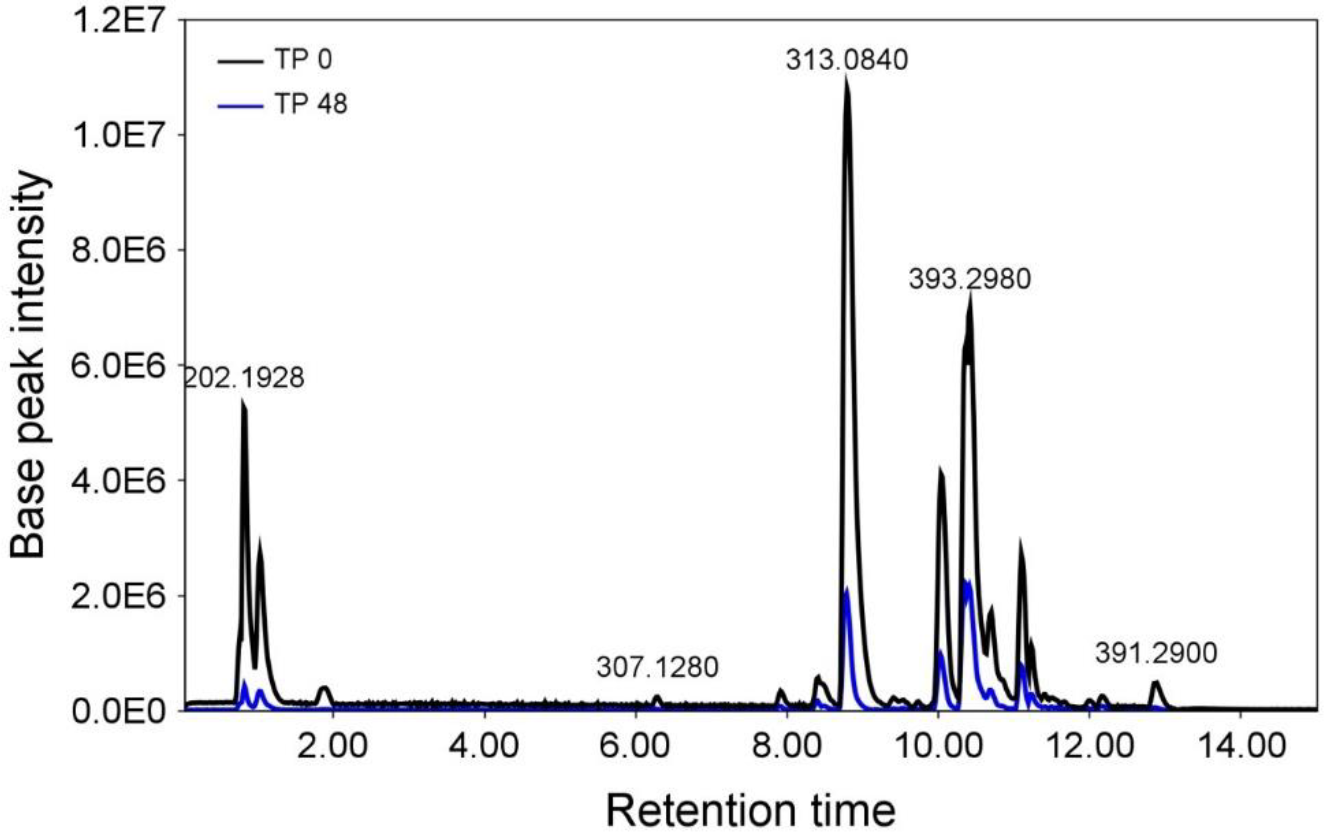
AFB_1_ degradation by pH 9 buffered medium does not show potential byproducts in the spectrum. The full mass spectra for time points (TP) 0 and 48 hours of incubation of AFB_1_ in pH 9 buffered medium, showing reduction of AFB_1_ (*m/z* 313) and no increased mass peak at the 48-hour time point. Retention times are in minutes. The corresponding *m/z* value is shown for each identified peak.

